# TGFβ restricts T cell function and bacterial control within the tuberculous granuloma

**DOI:** 10.1101/696534

**Authors:** Benjamin H Gern, Kristin N Adams, Courtney R Plumlee, Caleb R Stoltzfus, Laila Shehata, Kathleen Busman-Sahay, Scott G Hansen, Michael K Axthelm, Louis J Picker, Jacob D Estes, Kevin B Urdahl, Michael Y Gerner

## Abstract

Interferon gamma (IFNγ) produced by CD4 T cells is required for immune containment of *Mycobacterium tuberculosis* (Mtb) infection. Despite this, IFNγ plays a minor role in CD4 T cell-mediated immunity within the lung. In this study, we use a recently-developed murine model of physiologic Mtb infection coupled with advanced quantitative imaging to demonstrate that IFNγ production by Mtb-specific T cells is rapidly extinguished within the granuloma, but not in unaffected areas of the lung. This is mediated via localized immunosuppression through cell-intrinsic TGFβ signaling in effector T helper 1 cells within the granuloma, and blockade of TGFβ signaling in T cells results in improved immune cell function and decreased pulmonary bacterial burden. These findings uncover a potent immunosuppressive mechanism associated with Mtb infection and provide potential targets for host-directed therapy.

## INTRODUCTION

The tuberculous granuloma, an organized aggregate of macrophages and other immune cells, is the epicenter of *Mycobacterium tuberculosis* (Mtb) infection in the lung (Pagán and Ramakrishnan, 2015). For years, the granuloma was thought to be host-beneficial, walling off Mtb infection and providing protection against disseminated disease. More recently, however, there has been growing awareness of the role of the granuloma in pathogenesis. Pathogenic mycobacteria have been shown to promote macrophage aggregation and early granuloma formation to facilitate bacterial expansion and dissemination (Davis and Ramakrishnan, 2009; Volkman et al., 2004, 2010). Later during infection, mature granulomas provide a niche for Mtb persistence. It is commonly believed that granulomas provide an immunosuppressive milieu that restricts anti-Mtb immune effector function (Ernst, 2018), but the mechanistic basis of this purported inhibition is poorly understood. Elucidating pathways that impair effective immunity within granulomas could suggest therapeutic approaches that shorten and improve treatment of tuberculosis (TB).

CD4 T cells are critically important for immunity against Mtb (Cooper, 2009; Sileshi et al., 2013) and their protective capacity depends dually on their function and their migratory properties. IFNγ produced by CD4 T cells is a key contributor to protection (Flynn et al., 1993; Green et al., 2013). Although IFNγ accounts for almost all the protective capacity of CD4 T cells in the spleen, for unknown reasons, it is responsible for less than half of CD4-mediated immunity in the lung (Sakai et al., 2016). In addition to proper function, CD4 T cells must traffic to sites of infection and directly recognize antigen presented by infected cells to confer optimal protection (Srivastava and Ernst, 2013). In mice, we and others have shown that terminally-differentiated Mtb-specific CD4 T cells that are robust producers of IFNγ traffic poorly into the lung parenchyma and are not protective upon adoptive transfer, whereas less differentiated T cells that enter the parenchyma are protective (Moguche et al., 2015; Sakai et al., 2014). In non-human primates (NHP), T cells accumulate in the lymphocyte cuff surrounding the granuloma, but are present in lesser numbers in the central granuloma core where Mtb-infected cells reside. Inhibition of tryptophan metabolism has been shown to promote T cell re-localization to the center of the granuloma and decrease bacterial burdens (Gautam et al., 2017). These observations underscore the importance of T cell effector responses occurring in the right location, as well as reveal the likely presence of inhibitory mechanisms within the granuloma to modulate optimal T cell function.

Immune responses to Mtb infection are heterogeneous across different individuals, as well as among distinct granulomas within the same host. This is best demonstrated by recent findings in NHPs that each granuloma is initiated by a single bacterium, and these distinct lesions display variations in immune cell organization and infection outcomes (Marakalala et al., 2016), ranging from clearance to rapidly progressive infection (Lin et al., 2013). When coupled with the lack of genetic tools available in NHPs, this extensive heterogeneity makes it challenging to decipher the molecular mechanisms responsible for limiting cellular immunity to Mtb infection. On the other hand, while mice offer an array of experimental tools for mechanistic studies, they form poorly-organized immune cell aggregates that lack many features of human Mtb granulomas, at least using the standard model in which mice are infected via aerosolization with 50-100 CFU. Therefore, there is a need for more tractable experimental models coupled with advanced in situ imaging approaches to elucidate the composition, spatial organization, and function of T cells and other immune cells within granulomas that are associated with various Mtb infection outcomes.

We sought a model system with well-organized granuloma structures, but also one that was experimentally-tractable. In an effort to improve the murine TB model, we recently developed an ultra-low dose (ULD) infection model that uses a physiologic aerosolized dose of 1-3 CFU (Plumlee, unpublished data). Importantly, most mice infected with ULD Mtb develop a single, well-organized infectious lesion, founded by a single bacterium, which exhibits spatial segregation of infected macrophages and the surrounding lymphocytic cuff, thus recapitulating hallmark features of human Mtb granulomas.

Here, we use multiplex confocal imaging plus histocytometry (Gerner et al., 2012) to characterize the spatial organization of the CD4 T cell response during pulmonary ULD Mtb infection. Our findings reveal that CD4 T cell infiltration into granulomas is associated with partial TCR-mediated activation but poor production of IFNγ, even in models where T cells undergo optimized Th1 effector polarization. We find that abundant TGFβ signaling by immune cells within the granuloma core locally inhibits effector T cell function, and that conditional ablation of the TGFβ receptor (TGFβR) rescues T cell function and reduces bacterial burden. These studies identify an important pathway restricting T cell-mediated immunity in the Mtb granuloma and implicate TGFβ signaling as a potential target for host-directed therapy.

## RESULTS

### IFNγ production by CD4 T cells is diminished in the pulmonary Mtb granuloma

To characterize the spatial organization of CD4 T cell IFNγ production and antigen sensing within the pulmonary tuberculous granuloma, we used the murine ULD aerosol infection model, in which mice develop a solitary pulmonary granuloma. This granuloma is typically well-organized and recapitulates several key features of human Mtb granulomas, including spatial segregation of infected myeloid cells from CD4 T cells (analogous to the central core of infection with surrounding lymphocytic cuff), as well as presence of B cell aggregates, consistent with early tertiary lymphoid structure formation (Figs 1A, 1B) (Kahnert et al., 2007). This model provided an opportunity to investigate the localization of T cell antigen sensing and IFNγ production within tuberculous granulomas in an experimentally-tractable system.

**Figure 1:**
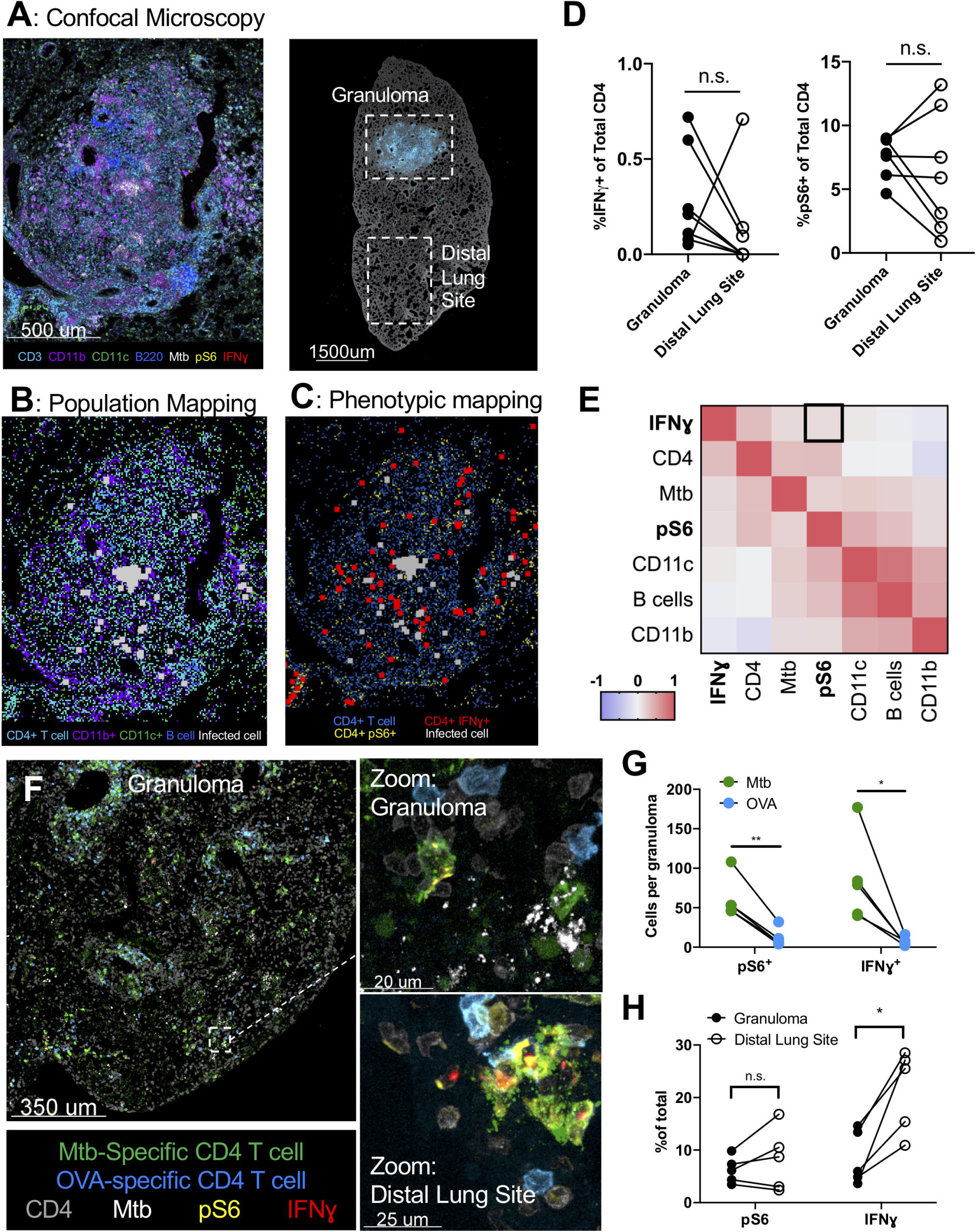
Despite antigen sensing, CD4 T cells produce minimal IFNγ within Mtb granulomas. <u>A-E: Wild-type mice were infected with ULD Mtb and lungs were harvested d34 p.i</u>. A) Representative 12 parameter confocal microscopy image of a granuloma from a 20um infected lung tissue section. B) Location of different cell types within the granuloma using histocytometry and positional mapping. Gating strategy is shown in Figure S1. C) Location of CD4 T cells that have produced IFNγ and recently sensed antigen in relation to Mtb. D) Image (left) showing whole lobe from another lung, demonstrating strategy for analysis for investigating granuloma and distal lung sites. Quantification (right) of IFNγ production and antigen-sensing (pS6^+^) in CD4 T cells as quantified by the relative location within the lung. E) Spatial correlation analysis of different cell types within local tissue neighborhoods. <u>F-G: Mice were infected with ULD Mtb and 34 days after infection received an adoptive transfer of 5×10^6^ Th1-polarized Mtb-specific and OVA specific cells. Lungs were harvested for microscopy 24 hours later.</u> F) Representative confocal microscopy image, showing Mtb (green) and OVA (blue) specific Th1-polarized CD4 T cells. G) Quantification of IFNγ and pS6 staining in Mtb vs OVA -specific transferred cells, as determined by histocytometry. H) Relative staining for pS6 and IFNγ in Mtb-specific CD4 T cells in different parts of the lung (granuloma vs distal lung site). Single-group comparisons by paired t test. Correlations and corresponding P values (E) by Pearson’s correlation test. *p < 0.05, ***p < 0.001, ****p < 0.0001. See also Figure S1.

To visualize the location of polyclonal T cell activation relative to Mtb-infected cells within these granulomas, we performed multiplex confocal microscopy on lung sections from mice infected with ULD Mtb 34 days prior. Sections were stained with a panel of antibodies recognizing various immune cell lineage-defining markers, as well as functional proteins, such as IFNγ and ribosomal protein phospho-S6 (pS6), which is downstream of TCR signaling, shown to be upregulated within one hour following TCR signaling in vitro in T cells (Sauer et al., 2008), and is downstream of antigen exposure in Mtb infected lungs (Delahaye et al., 2019). We also stained sections with an antibody to Mtb purified protein derivative (PPD) to identify Mtb-infected cells (Mehra et al., 2012), as fluorescent Mtb strains undergo plasmid loss over time and do not reliably delineate infected cells at later time points (not shown). Using histocytometry (Gerner et al., 2012), a method for quantitative image analysis that allows characterization of complex cell phenotypes while retaining information on cell positioning within tissues, we found that Mtb was harbored almost exclusively by CD11b^+^ myeloid cells located within the central core of the granuloma (Figs 1A, 1B). T cell infiltration was observed throughout the lung, but with greater overall density in and around the granuloma, with the infiltrating T cells spatially segregated from the core of infected cells. We next compared the functional properties of T cells within the granulomas versus those in the same lobe of the lung, but distal to the granulomatous site of infection (Figs 1D, S1A). The positioning of CD4 T cells that recently recognized antigen (pS6^+^) or produced IFNγ were visualized and quantified in relation to other cells types and Mtb (Figs 1B, 1C). Despite Mtb infection being contained solely within the granuloma, we observed no difference in the proportion of CD4 T cells that recently recognized antigen (5-10% pS6^+^) or produced IFNγ (<1%) within the granuloma compared to those in distal regions of the lung (Fig 1D). To obtain spatial granularity on how localized TCR signaling and effector function correlate across the granulomas, we used a virtual segmentation algorithm to subdivide the histocytometry datasets into individual 80μm radius neighborhoods (Fig S1B) (Stoltzfus, unpublished data), as this distance has been previously shown to be the maximum range of IFNγ signaling at sites of infection (Müller et al., 2012). We then investigated how different cell types and phenotypes were correlated within these 80um neighborhoods. This analysis revealed that neighborhoods with higher numbers of T cells that recently sensed TCR signals (pS6^+^) were only weakly correlated (R^2^=0.14, p<0.0001) with those having more IFNγ^+^ CD4 T cells (Fig 1E, S1D). These data suggest that the paucity of IFNγ production by effector CD4 T cells residing within the tuberculous granuloma is only partially explained by limited antigen recognition and that additional inhibitory mechanisms may be at play.

### Inhibition of IFNγ production by Mtb-specific CD4 T cells

Given that these findings were obtained from polyclonal CD4 T cells, which are comprised of both Mtb-specific and T cells with irrelevant specificities, as well as those with varying differentiation states, we next tested whether fully functional Mtb-specific Th1 effector CD4 T cells would demonstrate similar properties. To this end, we adoptively transferred 5×10^6^ Th1-polarized TCR-transgenic CD4 T cells (C7) that were specific for the Mtb antigen ESAT-6 (Gallegos et al., 2008), as well as TCR-transgenic CD4 T cells (OT-II) with an irrelevant specificity (OVA) (Barnden et al., 1998), into wild-type mice infected with ULD Mtb 34 days prior. Twenty hours after transfer, lungs were taken for microscopy and flow cytometry.

Although both Mtb-specific and control OVA-specific T cells showed evidence of trafficking into pulmonary granulomas within 20 hours after transfer (Fig 1F), Mtb-specific cells comprised the vast majority of transferred T cells that expressed pS6 and were IFNγ^+^ by histo- and flow cytometry (Fig 1G, S1E), suggesting in vivo Mtb-antigen recognition. Notably, the relative ratio of Mtb-specific vs. OVA-specific cell isolation was different with histocytometry as compared to flow cytometry (Fig S1F), which may indicate that the differences were underestimated in the flow-based assay due to differential loss of activated cells with dissociation-based methods (Borges da Silva et al., 2019; Steinert et al., 2015). Despite their prior capacity to robustly produce IFNγ upon in vitro peptide stimulation immediately prior to transfer (~20h before tissue harvest) (Fig S1G), we found diminished IFNγ production by Mtb-specific T cells located within the granuloma as compared to those located in the distal lung parenchyma (Fig 1H). This difference did not seem to reflect reduced antigen recognition within the granuloma, as pS6 expression was similar in both locations (Fig 1H), mirroring the pattern found in the polyclonal population (Fig 1D). Taken together, these results suggest that IFNγ production by Mtb-specific CD4 T cells is restricted locally by the granuloma microenvironment despite ongoing antigen recognition within these structures.

### TGFβ signaling in CD4 T cells in the Mtb-infected lung parenchyma

To identify potential suppressive mediators influencing T cell function within the Mtb-infected lung parenchyma, we sorted Mtb ESAT-6 specific CD4 T cells using MHCII tetramers and compared mRNA expression of vascular vs. parenchymal Mtb-specific T cells (as determined by intravascular labeling). We, as well as others, have previously shown that Mtb-specific CD4 T cells residing in the lung parenchyma exhibit distinct phenotypic and functional properties compared to T cells in the lung vasculature, including a diminished capacity to produce IFNγ (Moguche et al., 2015; Sallin et al., 2017). We found upregulation of a number of genes associated with T cell immunosuppression within the lung relative to the vasculature, including *TIGIT*, *CTLA4*, *Lag3*, *CCR8* and *HAVCR2* (encodes Tim3), and *SMAD3* (downstream of TGFβ) (Fig 2A). To better understand which inhibitory pathways had the greatest downstream effects on T cells, we used gene set enrichment analysis to identify transcriptional signatures associated with immunosuppression. Within the lung, we found upregulation of IL-2, hypoxia and glycolysis pathways. Importantly, we also observed strong enrichment in target genes downstream of TGFβ signaling within the lung parenchymal Mtb-specific T cells as compared to their counterparts in the vasculature (Figs 2B, S2C). TGFβ has been previously observed to restrict T cell proliferation and differentiation and can directly inhibit IFNγ production (Oh and Li, 2013). Furthermore, an Mtb cell wall component (ManLam) has been shown to potently induce TGFβ production in macrophages (Dahl et al., 1996), and TGFβ production has been shown to increase dramatically in the lungs of Mtb-infected mice (Rook et al., 2007). TGFβ has also been previously detected in bronchoalveolar lavage specimens from patients with active pulmonary TB (Bonecini-Almeida et al., 2004), and lung TGFβ levels decrease with antibiotic treatment in NHPs (DiFazio et al., 2016). Thus, TGFβ signaling emerged as a strong candidate as a potential mediator of local CD4 T cell suppression in the granuloma.

**Figure 2:**
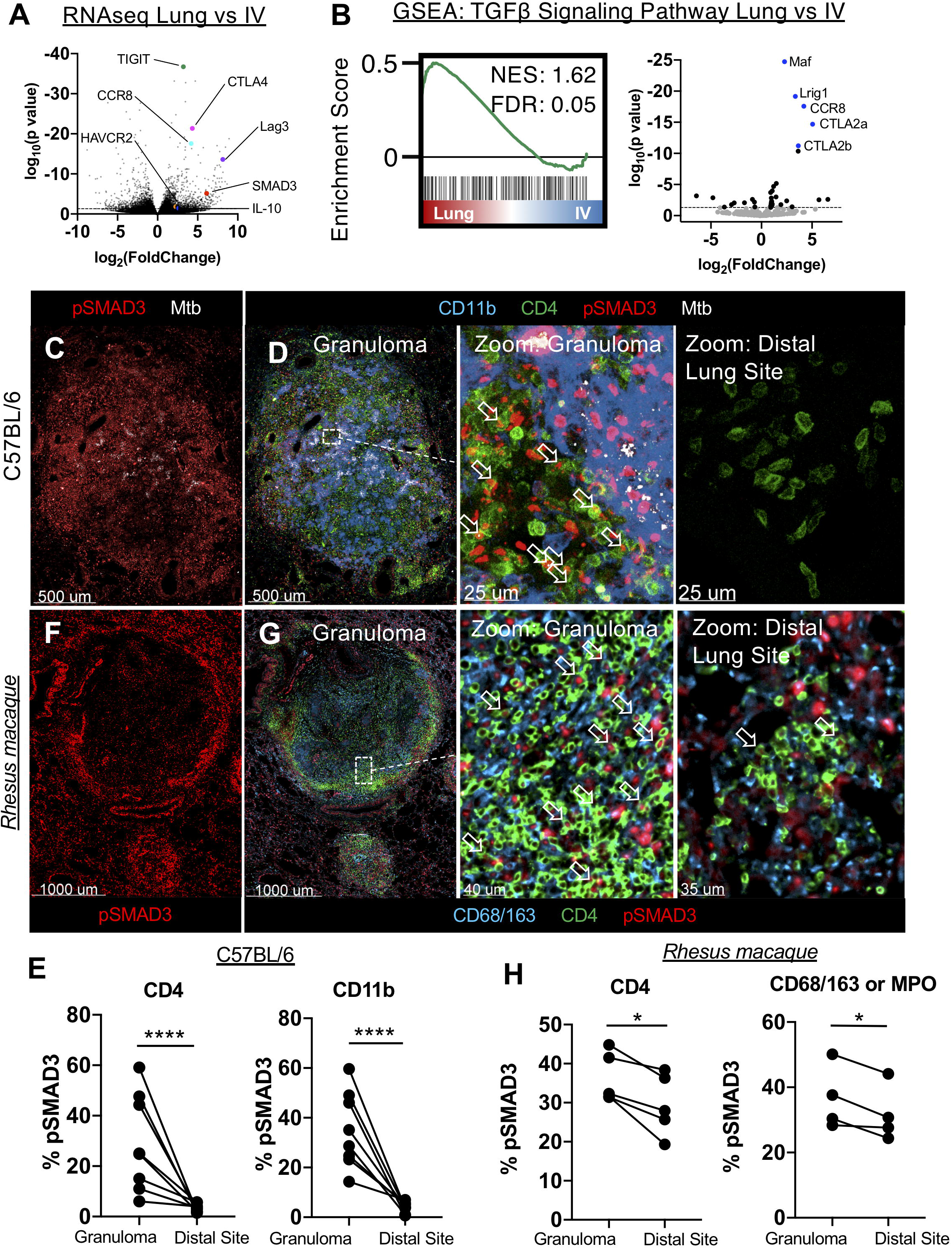
TGFβ signaling by immune cells within the granuloma. <u>A-B: WT mice were infected with 50 CFU Mtb, then lungs taken at 28 days p.i. for transcriptome analysis of ESAT-6 specific CD4 T cells in the vasculature vs lung parenchyma.</u> A) Volcano plot of differentially expressed genes by Mtb-specific T cells within the parenchyma vs intravascular space. Highlighted genes are associated with immunosuppressive signals on T cells. B) GSEA analysis of genes downstream of TGFβ signaling, ESAT-6 specific CD4 T cells in the parenchyma (Lung), versus vasculature (IV). <u>(C-E) Mice were infected with ULD Mtb and lungs were harvested at d35 p.i. for analysis.</u> C) Representative confocal image of pSMAD3 staining within the granuloma. D) Zoom-in images demonstrating pSMAD3 staining within T and myeloid cells near Mtb (white) infected cells (left), or at a distal lung site (right). E) Histocytometry analysis of pSMAD3 staining in CD4 T cells and CD11b+ myeloid cells in infected mouse lungs. F) Representative confocal image of an NHP granuloma, showing pSMAD3 staining. G) Zoom-in images showing pSMAD3 signal in relation to CD4 and CD68/163. H) Histocytometry of NHP lungs, showing pSMAD3 staining in CD4^+^ and CD68/163^+^ cells. Single-group comparisons by paired t test. *p < 0.05, ***p < 0.001, ****p < 0.0001. Data are representative of two independent experiments. FDRs were calculated using the Benjamini-Hochberg method and p values were calculated using the Mann-Whitney U test. See also Figure S2.

To examine if there was active TGFβ signaling occurring within Mtb-infected lungs, we examined SMAD3 phosphorylation (pSMAD3), downstream of TGFβ signaling (Derynck and Zhang, 2003), in d34-infected lungs by microscopy. We observed robust pSMAD3 staining within the granulomas in both CD4 T cells (6-59%) and CD11b^+^ myeloid cells (14-59%), though not in unaffected areas of the same lobe (Figs 2C-E). Histocytometry showed predominant localization of pSMAD3-positive T cells and myeloid cells within the granuloma, and not in the unaffected, distal areas of the same lobe (Fig 2E), suggesting highly localized and ongoing TGFβ signaling by T cells and myeloid cells within the pulmonary Mtb granulomas. We next assessed pSMAD3 expression in pulmonary Mtb granulomas in rhesus macaques that had been infected with 4-8 CFU via bronchoscope instillation 63-89 days prior. Similar to our data in mice, we found that there was evidence of strong pSMAD3 signaling within T cells (30-45%), as well as and macrophages and neutrophils (28-50%) within the granuloma, particularly in the lymphocytic cuff, in both non-necrotic (Figs 2F,2G) and necrotic granulomas (Figs S2D, S2E). Although rhesus macaques exhibited higher baseline pSMAD3 expression in distal areas of the lung compared to mice, which could be reflective of species, animal age, infectious dose, time point, or microbiome differences, pSMAD3 expression was significantly increased for CD4 T cells and macrophages/neutrophils residing within the granuloma compared to distally (Fig 2H). Together these data indicate that T cells infiltrating pulmonary Mtb granulomas experience localized TGFβ signaling in both mice and non-human primates.

### Lack of TGFβRII on T cells results in decreased CFUs and increased IFNγ production

To test whether TGFβ actively suppresses T cell function and restricts immunity within the granuloma we used *dLck*-cre *TGFβR2*^f/f^ mice (TGFβR.KO), which lack TGFβR on T cells but express the receptor normally on other immune cell types (Zhang and Bevan, 2012). We first infected these animals with a conventional dose of aerosolized Mtb (50 CFU) in order to ascertain the role of TGFβ in modulation of immune responses and infectious outcome using an infection model with well-established bacterial load kinetics. We found that TGFβR.KO mice exhibited a significantly reduced bacterial burden at d28 p.i., suggesting that intrinsic TGFβ signaling in T cells restricts anti-Mtb immunity (Fig 3A). Despite this decreased bacterial burden, which likely corresponds to reduced antigen abundance, we found that TGFβR.KO mice contained a trend towards a higher proportion of IFNγ-producing CD4 T cells specific for the Mtb antigen ESAT-6 compared to their wild-type counterparts (Fig 3B). Indeed, in mice with a similar bacterial burden there was a marked increase in IFNγ in TGFβR.KO mice (Fig 3C). Together this suggests that alleviating TGFβR signals on T cells results in increased IFNγ.

**Figure 3:**
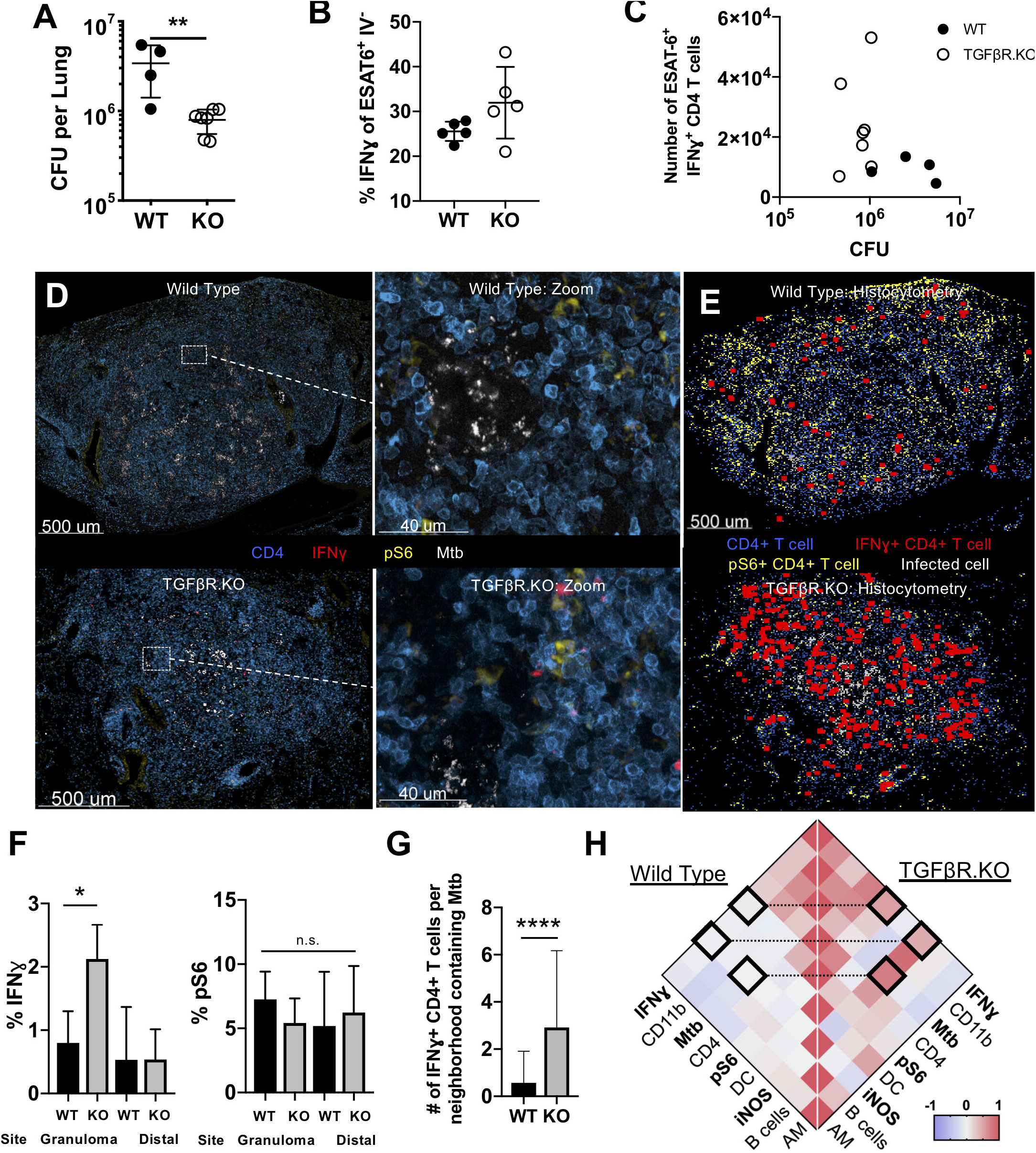
Lack of TGFβRII on T cells results in decreased CFUs and increased IFNγ production. <u>A-B) TGFβR.KO (KO) and WT mice were infected with 100 CFU of Mtb and lungs were taken for analysis d30 p.i</u>. A) Mtb lung burden was analyzed in WT and KO animals. B) Flow cytometry of IFNγ production on ESAT-6 specific CD3^+^CD4^+^ T cells. C) Number of IFNγ^+^ ESAT-6 specific CD4 T cells compared against bacterial burden. <u>(D-H) WT mouse and in TGFβR.KO mice were infected with ULD Mtb and lungs were harvested d35 p.i</u>. D) Representative confocal images of granulomas from WT (top) and TGFβR.KO (bottom) mice, with zoom-ins. E) Histocytometry positional analysis demonstrating the location of pS6^+^ or IFNγ+ cells in WT and TGFβR.KO lung granulomas. F) Histocytometry quantitation of CD4 cells positive for pS6 and IFNγ in the granuloma vs a distal lung site. G) Number of IFNγ^+^ CD4 T cells per local tissue neighborhood containing Mtb-infected cells was quantitated in WT vs. TGFβR.KO infected lungs. H) Comparative analysis of cell-cell correlations across local neighborhoods in WT (left) and TGFβR.KO (right) infected lungs is shown. Single-group comparisons (A,B,F,G) by unpaired t test. Correlations and corresponding P values (H) by Pearson’s correlation test. Data are representative of two independent experiments. Data are presented as mean ± SD. *p < 0.05, ***p < 0.001, ****p < 0.0001. See also Figure S3.

To further delineate how the absence of TGFβ signaling influences localized effector T cell functionality within well-organized granulomas, we infected TGFβR.KO and wild-type mice with ULD Mtb and isolated lungs for microscopy 34 days later for quantitative imaging. We found that TGFβR.KO granulomas exhibited a pronounced increase in IFNγ production adjacent to the areas of Mtb infection as compared to WT mice (Figs 3D,3E). In particular, CD4 T cells from TGFβR.KO mice made significantly more IFNγ directly inside the granuloma, while there was no significant difference within distal lung sites (Fig 3F). No differences in pS6 staining of CD4 T cells were observed between TGFβR.KO and WT mice (Fig 3F), suggesting that antigen presentation and TCR-signaling were not significantly increased by TGFβ. These results suggest that TGFβ potently suppresses effector T cell function within the granuloma, but not the remainder of the lung.

To explore how localized IFNγ production by T cells influences immune responses within the granulomas of TGFβR.KO animals, we again analyzed the images by virtually segmenting the histocytometry data and performing cell-cell correlation analysis (Fig S1B). We found that IFNγ+ CD4+ T cells were more frequent in neighborhoods containing Mtb in TGFβR.KO mice (2.9 per 80um neighborhood, SD 3.3) than in WT mice (0.5 per 80um neighborhood, SD 1.4) (Fig 3G). Furthermore, neighborhoods containing IFNγ+ CD4+ T cells were correlated with those containing T cells that had undergone antigen-sensing in TGFβR.KO (R^2^ = 0.53, p<0.0001) but not WT (R^2^= 0.03, p=0.06) mice (Figs 3H, S3B), suggesting restoration of localized effector T cell function in the absence of TGFβ signaling. Given that the production of IFNγ by T cells may be short-lived, and would be challenging to detect after cell-mediated secretion, we also examined myeloid cell iNOS production (Fig S3C), a biomarker reflecting cellular memory of IFNγ sensing as well as a molecule directly involved in mycobactericidal activity (Braverman and Stanley, 2017). The number of iNOS^+^ CD11b^+^ cells was directly associated with Mtb-containing neighborhoods in TGFβR.KO mice (R^2^ = 0.76, p<0.0001) as compared to WT animals (R^2^ = −0.01, p=0.54), as well as more associated with IFNγ+ CD4+ T cells in TGFβR.KO mice (R^2^ = 0.44, p<0.001) but not WT animals (R^2^ = −0.03, p=0.05) (Fig 3H, S3C), both suggesting enhanced coupling of localized immune cell function with myeloid cell activation in the absence of TGFβ sensing. Together, these data indicate that inhibition of TGFβ signaling in T cells results in marked enhancement in granuloma-localized effector T cell function, increased downstream myeloid cell activation, and decreased bacterial burden.

### TGFβ inhibits effector function of Th1 cells in a cell-intrinsic manner

To further investigate the intrinsic role of TGFβ signaling in restricting IFNγ production by CD4 T cells, we generated mixed bone marrow chimera mice containing both WT and TGFβR.KO T cells and infected these animals with ULD Mtb. Thirty days later, we compared the phenotype and function of WT and TGFβR.KO CD4 T cells isolated from the same lungs, and therefore subjected to the same bacterial burden and inflammatory milieu. We found a higher percentage of IFNγ^+^ Mtb ESAT-6-specific CD4 T cells among TGFβR.KO than their WT counterparts, and those that did produce IFNγ produced a greater amount (Figs 4A, S4B). This confirmed that TGFβ was acting in a T cell intrinsic manner and validated our observation (Fig 3C) that TGFβR.KO cells had higher IFNγ production than WT cells, when exposed to similar bacterial loads. Although WT and TGFβR.KO cells were equally represented amongst naïve (CD44 low) CD4 T cells in the chimeras, ESAT-6 tetramer-binding cells were higher in the TGFβR.KO cells (Fig 4B). Furthermore, a higher proportion of the ESAT-6-specific TGFβR.KO T cells in the lung parenchyma expressed KLRG1 (Fig. 4C), a marker of terminally differentiated CD4 T cells,, which are almost exclusively found in the vasculature in WT mice (Moguche et al., 2015; Sakai et al., 2014). Thus, T cell intrinsic TGFβ-signaling has a wide range of downstream effects, shaping the abundance, differentiation, and function of Mtb-specific CD4 T cells in the infected lung.

**Figure 4:**
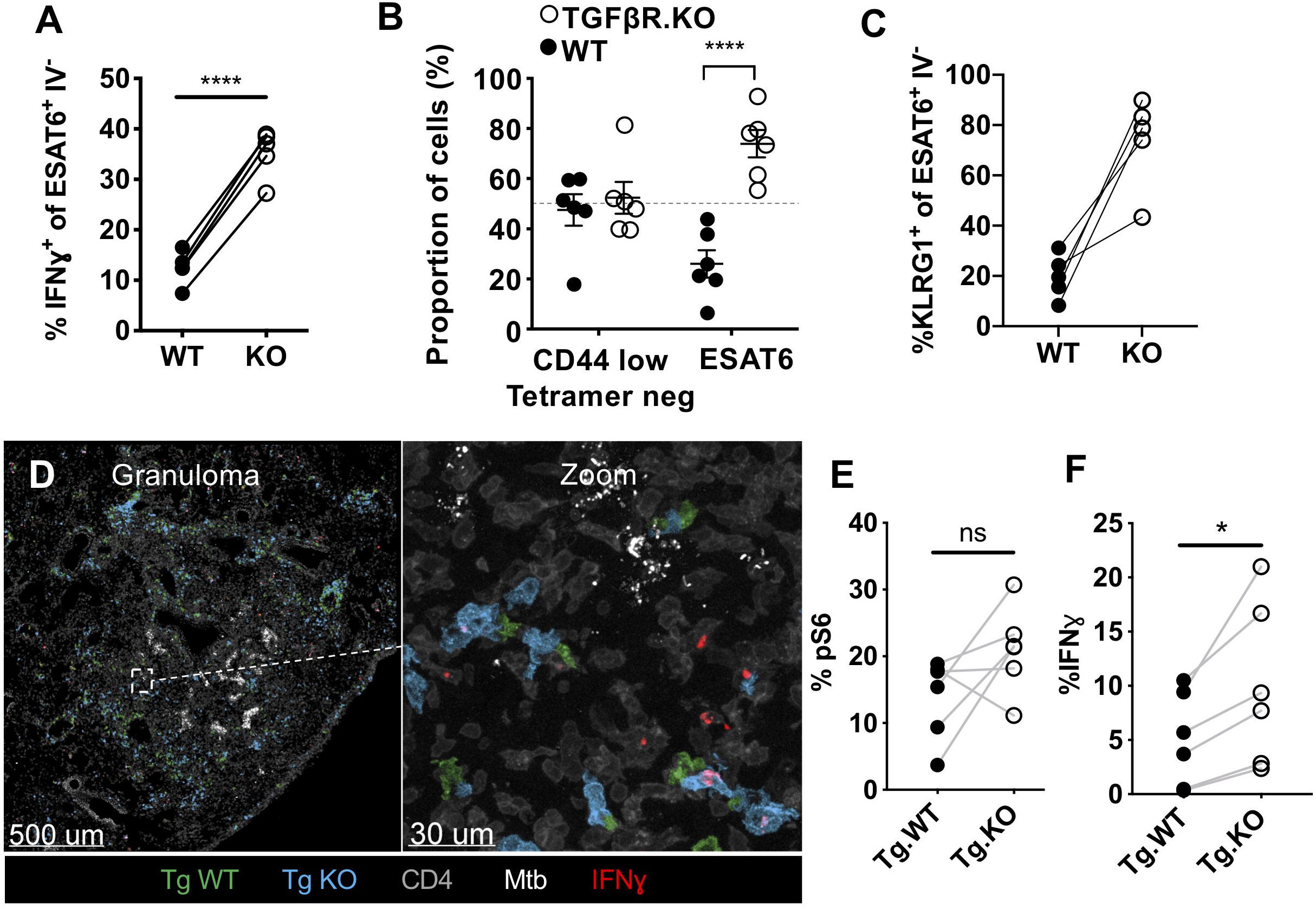
TGFβ inhibits effector function of Th1 cells in a cell-intrinsic manner. <u>(A-B) Mixed WT:TGFβR.KO bone marrow chimeric mice were infected with 100 CFUs of Mtb by aerosol.</u> A) IFNγ production by parenchymal ESAT-6-specific cells after ex vivo restimulation was analyzed. Lines connect WT and KO cells within the same chimeric animals. B) Relative frequencies of detected ESAT-6 specific WT and TGFβR.KO cells are shown. C) Percentage of parenchymal ESAT-6 specific CD4 T cells that express KLRG1 in WT vs TGFβR.KO mice is quantified. Lines connect WT and KO cells within the same animals. <u>(D-F) Mice were infected with ULD Mtb and d34 p.i. received an adoptive transfer of WT (Tg.WT) and TGFβR.KO (Tg.KO) Mtb-specific Th1-polarized CD4 T cells. Lungs harvested 20 hours after transfer.</u> D) Representative section (left) demonstrating presence of Tg.WT (green) and Tg.KO (blue) cells within the granuloma. Zoom-in panel (right) showing localization of both cell types proximal to Mtb-infected cells, as well as demonstrating IFNγ staining. Histocytometry quantification of E) pS6 and F) IFNγ staining in WT vs KO transferred cells within the granuloma. Single-group comparisons by paired t test. Data are representative of two independent experiments. Data are presented as mean ± SD. *p < 0.05, ***p < 0.001, ****p < 0.0001. See also Figure S4.

To elucidate whether cell-intrinsic TGFβ signaling is capable of directly suppressing the function of fully differentiated effector CD4 T cells at the granuloma site, we generated mice with Mtb-specific TCR-transgenic T cells lacking TGFβR (Tg.KO). Th1 effector cells from Tg.KO and control Mtb-specific WT (Tg.WT) CD4 T cells were generated in vitro and co-transferred into mice infected with ULD Mtb 34 days prior. Importantly, the lack of TGFβR did not seem to impact Th1 polarization, as Tg.KO and Tg.WT T cells made similar amounts of IFNγ cells prior to transfer (Fig S4C). Lungs were isolated for microscopy 20 hours later. We found that both Tg.WT and Tg.KO Th1 T cells were able to infiltrate the granulomas (Fig 4D) and displayed similar levels of TCR signaling, as identified by pS6 staining (Fig 4E). Importantly, we found ~2.5 fold more Tg.KO T cells than Tg.WT cells within the granulomas (Fig 4F), suggesting enhanced effector function by T cells that are unable to receive TGFβ-mediated signals within the granuloma.

## DISCUSSION

IFNγ production by CD4 T cells is required for immune control of Mtb infection (Green et al., 2013). Despite this, IFNγ has been shown to account for only 30% of CD4-mediated immunity in the lung (Sakai et al., 2016). Here we use quantitative imaging to show that little IFNγ is produced by CD4 T cells within Mtb-infected granulomas, and that fully differentiated Mtb-specific Th1 effector T cells rapidly lose the ability to produce IFNγ within the pulmonary granuloma, as compared to distal areas of unaffected lung. This is despite ongoing antigen recognition by CD4 T cells both within the granuloma and in distal lung areas. We further find that effector T cell function was actively modulated by localized TGFβ within the granuloma, and that genetic manipulation of TGFβ signaling on T cells restores T cell function and limits bacterial burden in the infected lungs. Of note, in our transcriptional analysis, parenchymal Mtb-specific T cells had higher levels of *SMAD3* transcription, suggesting that they are more capable of responding to TGFβ signals, which involves phosphorylation of SMAD3. These findings provide a mechanistic explanation why effects of IFNγ are typically limited within the lung, as its production by T cells is potently inhibited via localized TGFβ-mediated immunosuppression.

Previous studies have shown elevated TGFβ levels in the lungs of Mtb-infected mice, NHPs and humans (Bonecini-Almeida et al., 2004; DiFazio et al., 2016; Rook et al., 2007). Our data now show that in both mice and NHPs T cells and myeloid cells experience high levels of TGFβ signaling within the pulmonary Mtb granuloma, and less within the rest of the lung. TGFβ signaling in turn rapidly suppresses effector T cell function even in fully differentiated Mtb-specific Th1 CD4 T cells, and ablating TGFβ signaling specifically on T cells rescues their functionality in a cell-intrinsic manner. Additional effects of TGFβ on T cell proliferation and altered differentiation/maturation also likely play a role in limiting protective immunity, and further studies are necessary to delineate these distinct processes.

Mtb is known to have a multitude of different strategies to evade the immune system (Ernst, 2018). We have shown that the neutralization of only one of these mechanisms on a single cell type results in a significantly decreased bacterial burden, despite the continued presence of additional intact mechanisms of immune suppression (including the effect of TGFβ on other cell types). In fact, here we demonstrate that TGFβ is sensed by CD11b^+^ cells within the granuloma, and it has been previously shown that TGFβ is an important factor in suppressing macrophage immunity to Mtb (Jayaswal et al., 2010). This underscores the role of TGFβ as an inhibitory mechanism of both T cell and myeloid cell function within Mtb granulomas, as well as suggests a potential application for TGFβ inhibitors in the treatment of Mtb infection. In support of this idea, pharmacologic inhibition of TGFβ signaling, initiated 10 days after aerosol Mtb infection, has been shown to prevent pulmonary granuloma formation and to reduce lung bacterial burdens (Jayaswal et al., 2010). Although these findings are promising, in clinical practice it would not be feasible to initiate treatment at such an early infection stage. In another study, however, aerosolized intrapulmonary delivery of siRNA targeting TGFβ was used to treat mice after chronic Mtb infection had been established and granulomas were already formed (Rosas-Taraco et al., 2011). Despite only achieving a 25% reduction in pulmonary TGFβ levels, this treatment modestly reduced the lung bacterial burden and also limited pulmonary fibrosis and immunopathology, raising the possibility that more effective inhibition could provide even greater benefit. While these studies did not resolve individual effects of TGFβ on specific cell types, together with our findings, they directly support the notion that therapeutic blockade of the TGFβ pathway during chronic disease can enhance T cell-mediated immunity directly at the site of infection with minimal off-target effects. Besides targeting T cells, global TGFβ blockade could provide additional therapeutic effects, including promoting anti-microbial function in macrophages and other cell types, as well as reducing fibrosis, which could improve pulmonary function (Walton et al., 2017).

An additional suppressive mechanism used by Mtb is the ability to limit MHCII antigen presentation within infected cells. Our observation that some pS6^+^ T cells reside adjacent to Mtb-infected macrophages in the granuloma core indicates that this inhibition is not complete. However, the fact that the proportion of pS6^+^ T cells in the granuloma core is similar to that in uninfected regions of the lung suggests that antigen-presentation by infected cells is impaired to some degree. We were surprised to find evidence of recent antigen recognition by T cells located throughout the lung. This TCR stimulation is likely due, at least in part, to recognition of exported antigen by Mtb to regions of the lung distal to infection, a possible decoy strategy that has been suggested to direct T cell responses to where they are not needed (Baena and Porcelli, 2009). Although such antigen-export has been shown to occur (uninfected cells have been shown to be better ex vivo antigen-presenters than infected cells isolated from the same Mtb-infected lungs) (Srivastava and Ernst, 2014; Srivastava et al., 2016), the location of this “decoy” T cell activation within the lung has not previously been examined. Alternatively, it is possible that some of the pS6^+^ T cells in uninfected regions of the lung represent cells that were activated within the lymph node in the prior 24 hours (the duration of pS6 expression after TCR engagement) (Sauer et al., 2008), and only recently immigrated to the lung. However, in our adoptive transfer experiments, we observed similar pS6 staining in Mtb-specific CD4 T cells distal to the granuloma 20 hours after transfer, which would provide a very limited window of opportunity for efficient trafficking to draining lymph nodes, local activation, and subsequent recirculation back into the lung.

What implications do our findings have about the role of CD4 T cells and IFNγ in Mtb infection? In our studies, mice with T cells lacking the TGFβ receptor have increased production of IFNγ and a decreased bacterial burden. On the surface this seems at odds with published findings that in PD-1 knockout mice (Lazar-Molnar et al., 2010), and patients who received PD-1 or PD-L1 blockade (Barber et al., 2019), there was an increased IFNγ response observed in concert with more severe disease outcomes. We speculate that the location of IFNγ production may help to reconcile these results. In our system, the increase in IFNγ production after TGFβ-blockade is greatest near Mtb-infected cells within the granuloma, while the above studies measured an increase in systemic IFNγ levels, which may not be reflective of granuloma-restricted processes, and could also trigger immune pathology. It is possible that checkpoint blockade alone is not sufficient to overcome TGFβ-mediated suppression within the granuloma. If so, combination immunotherapy may provide synergy. Immunomodulatory medications targeting the inhibition of TGFβ are currently under development for oncologic indications and have been demonstrated to be safe for administration to humans and are currently in phase III clinical trials (de Gramont et al., 2017). Our studies offer evidence to suggest a potential therapeutic role for TGFβ blockade during Mtb infection, which warrants further exploration in future studies.

## Acknowledgements

We thank Sara Cohen for comments on the manuscript and Sarah Stanley for helpful discussions. This study was supported by the NIH grants 1R01AI134246 (K.B.U.), 1R01AI076327 (K.B.U.), U19AI135976 (K.B.U.), R01AI134713 (M.Y.G.), R21AI142667 (M.Y.G.), and T32HD007233 (B.H.G.).

## Author Contributions

Conceptualization, B.H.G., K.N.A., K.B.U. and M.Y.G.; Formal analysis, B.H.G., K.N.A., C.R.S., K.B.U. and M.Y.G.; Funding acquisition, B.H.G., K.B.U. and M.Y.G.; Investigation, B.H.G., K.N.A., C.R.P., C.R.S., L.S., K.B., S.G.H., M.K.A., L.J.P., J.D.E., K.B.U., and M.Y.G.; Methodology, B.H.G., K.N.A., L.J.P., J.D.E., K.B.U. and M.Y.G., Project administration, K.B.U. and M.Y.G.; Resources, K.B., S.G.H., M.K.A., L.J.P., J.D.E.; Software, C.R.P; Supervision, K.B.U. and M.Y.G.; Validation, B.H.G., K.N.A., K.B.U. and M.Y.G.; Visualization, B.H.G., K.B.U. and M.Y.G.; Writing – original draft, B.H.G., K.B.U. and M.Y.G.; Writing – review & editing, B.H.G., K.N.A., C.R.P, J.D.E, K.B.U. and M.Y.G.

## Declaration of Interests

The authors declare no competing interests.

## Methods

### Mice

C57BL/6 and TCRβ^−/−^δ^−/−^ mice were purchased from Jackson Laboratories (Bar Harbor, ME). ESAT-6 TCR Tg (C7) mice were provided by Dr. Eric Pamer (Memorial Sloan Kettering Cancer Center, New York, NY) and have been described previously (Gallegos et al., 2008). OTII mice were obtained from Taconic Laboratories (Rensselaer, NY). *DLck*-cre *TGFβR2*^f/f^ mice were a generous gift from Dr. Michael Bevan (University of Washington). All mice were housed in specific pathogen-free conditions at Seattle Children’s Research Institute (SCRI). Experiments were performed in compliance with the SCRI Animal Care and Use Committee. Both male and female mice between the ages of 8-12 weeks were used.

### Generation of mixed bone marrow chimera mice

WT CD45.2 mice were irradiated with 1000 rads and reconstituted with a 3:1 mixture (to account for differences in reconstitution of CD45.1 TGFβR.KO:CD90.1 WT bone marrow.

### Aerosol infections

All infections were done with a stock of Mtb H37Rv, as previously described (Urdahl et al., 2003). To perform conventional dose aerosol infections, mice were enclosed in a Glas-Col aerosol infection chamber, and 50-100 CFU were deposited directly into their lungs. In order to confirm the infectious dose, two mice in each infection were immediately sacrificed and their lung homogenates plated onto 7H10 plates for CFU enumeration. To perform ULD aerosol infections, mice were enclosed in a Glas-Col aerosol infection chamber, and 1-3 CFU were deposited directly into their lungs (Plumlee et al., 2018).

### Th1 polarization and adoptive transfers

CD4 T cells from ESAT-6-specific (C7) CD90.1+ and OVA-specific (OTII) CD45.1+ TCR transgenic mice were negatively enriched from spleens using EasySep magnetic microbeads (STEMCELL). T cells were Th1 polarized by culturing 1.6 × 10^6^ transgenic T cells with 8.3 × 10^6^ irradiated splenocytes from TCRβ^−/−^δ^−/−^ mice per 2ml well. 5 μg/ml of ESAT-6 or OVA peptide, 10 ng/ml IL-12, and 10 μg/ml of anti–IL-4 antibody (R&D Systems) were added to RP10 media at day 0. On day 3, cells were split 2:1, and 10 ng/ml IL-12 added (R&D Systems). On day 5, Th1 cells were intravenously injected into C57BL/6 CD45.2+ mice infected with Mtb 35 days prior.

### Lung Single Cell Suspensions

At the indicated times post-infection, mouse lungs were removed and gently homogenized in HEPES buffer containing Liberase Blendzyme 3 (70 mg/ml; Roche) and DNaseI (30 mg/ml; Sigma-Aldrich) using a gentleMACS dissociator (Miltenyi Biotec). The lungs were then incubated for 30 min at 37°C and again thoroughly homogenized with the gentleMACS. Homogenates were then filtered through a 70 um cell strainer and RBC lysed with RBC lysing buffer (Thermo), then resuspended in FACS buffer (PBS containing 2.5% FBS and 0.1% NaN_3_).

### Antibody and MHCII Tetramer Staining

MHCII tetramers containing amino acids 4-17 of Mtb ESAT-6 (made in our lab using a construct generated by Dr. Marc Jenkins) were used to detect Mtb-specific CD4 T cells. Single-cell suspensions were stained at saturating concentrations with the tetramers and incubated at room temperature for 1 hour. For intracellular cytokine staining (ICS), tissues were processed in the presence of Brefeldin A (10 mg/ml; Sigma-Aldrich). All surface staining was done at room temperature for 60 minutes. After tetramer and/or surface staining, the cells were fixed, permeabilized, and stained with antibodies against intracellular targets at 4°C for 30 minutes using eBioscience fixation/permeabilization and permeabilization buffers. For experiments utilizing intravascular antibody to identify intravascular T cells, mice were given an intravenous injection in the retroorbital sinus of 0.25μg of CD4-PE or CD45.1-APC, or CD45.2 -APC antibody 10 min prior to sacrificing, as previously described (Sakai et al., 2014).

### Sorting, RNAseq, and GSEA

CD4 T cells were negatively enriched to >95% purity from fleshly isolated lungs (taken from mice receiving an intravascular label, as above) using Miltenyi Biotec magnetic microbeads and subsequent column purification according to the manufacturer’s protocol. The negatively enriched cells were stained with anti-mouse PD-1 and anti-mouse KLRG1 antibodies (BioLegend), as well as ESAT-6 tetramer (as above) presence of 1μg/ml Cyclosporine A (R&D Systems) to prevent tetramer-mediated activation of T cells. The cells were then sorted on a cell sorter (FACSAria; BD Bioscience) directly into a buffer containing TRIzol (Invitrogen). Samples were shipped to Expression Analysis (Durham, NC), RNA was isolated and amplified linearly followed by RNA sequencing. RNASeq data were analyzed at the Institute for Systems Biology (Seattle, WA). The Spliced Transcripts Alignment to a Reference (STAR) tool was used to map the RNASeq reads to University of California Santa Cruz (UCSC) mouse genome. Aligned reads were sorted and mapped to transcripts and counts were output into a matrix using HTseq-count. Quality control of samples was conducted by ensuring biological replicates clustered together more closely than samples from other test conditions using Principal Component Analysis (PCA). Differentially expressed genes were discovered by inputting the counts matrix into the DESeq2 package in R with Benjamini-Hochberg corrected p-values less than or equal to 0.05 and fold-change greater than or equal to 2. We used Gene Set Enrichment Analysis (GSEA, Broad Institute) to analyze the enrichment dataset using the Molecular Signatures Database (MSigDB). GSEA identified sets of genes differentially expressed between comparison groups that were over-represented at the top or bottom of the ranked set of genes. Data will be deposited in GEO upon publication.

### Immunofluorescent staining and imaging of rhesus macaques

All rhesus macaques (RM) were infected with low (4-8 CFU) or high (30-100 CFU) dose Mtb Erdman strain (BEI Resources) into the right caudal lobe using a bronchoscopy. Lung tissues were collected from animals according to BSL-3 necropsy procedures. Tissues were dissected into ~ 20mm^2^ × 5mm thick sections, placed in pre-labeled cassettes and immersed in freshly prepared 4% paraformaldehyde for 24 hours at room temperature in the ABSL-3 necropsy laboratory. This fixation protocol resulted in complete inactivation of Mtb, as no bacterial growth was noted after 8 weeks of culture with homogenized fixed tissue. After 24 hours post paraformaldehyde fixation the cassettes were transferred to 70% EtOH for an additional 24-72 hours at ~4-8oC in the ABSL-3 necropsy laboratory prior to processing and paraffin embedding. Immunofluorescent staining was performed as previously published (Mudd, Nature Communications, 2018) on formaldehyde-fixed, paraffin-embedded right caudal lung tissue collected at necropsy from Mtb infected RM. In brief, HIER was performed on deparaffinized slides with 0.1% citraconic anhydride (125318; Sigma-Aldrich) buffer in the absence of enzymatic retrieval, followed by rounds of sequential antibody staining, detection, and stripping. Antibody detection was performed using species specific polymer HRP-conjugated systems (GBI Labs) coupled with tyramide signal amplification (TSA) reactions using Alexa Fluor tyramide reagents (Invitrogen). Antibody stripping was performed by heating slides in 0.1% citraconic anhydride or citrate pH6 (B05C-100B; GBI Labs) buffers for 15 minutes at 95-99^°^C on a hot plate. Whole slide imaging was performed on a Zeiss Axio Scan.Z1 using a 20x objective.

### Antibodies

The following antibodies were used for staining mouse tissue sections for imaging or isolated cells for flow cytometry: CD11c (clone HL3; BD), CD11b (clone M1/70; BD), CD45.1 (clone A20; Thermo Fisher), Ly6G (clone IA8, BioLegend), CD45.2 (clone 104; BioLegend), MHCII (clone M5/114.15.2; BioLegend), CD3 (clone 17A2; BioLegend), CD62L (clone Mel-14; BD Biosciences), CD8 (clone 53-6.7; BioLegend), IFNγ (clone XMG1.2; eBiosciences), CD45.2 (clone 104; BioLegend), pS6 (clone 2F9; Cell Signaling), CD4 (clone RM4-4; BioLegend), CD4 (clone RM4-5; BioLegend), CD8a (clone 53-6.7; BioLegend); CD44 (clone IM7; BioLegend), KLRG1 (clone 2F1; BioLegend), PD-1 (clone RMP1-30; BioLegend), B220 (clone RA3-6B2; ThermoFisher), NOS2 (clone C-11; Santa Cruz Biotechnology), Anti-purified protein derivative (polyclonal ab905; Abcam), CD90.1 (clone HIS51; ThermoFisher), pSMAD3 (clone EP823Y; Abcam.

The following antibodies were used in Rhesus macaque experiments for staining sections for imaging: pSMAD3 (clone EP823Y; Abcam), myeloperoxidase (A0398; Dako), CD68 (clone KP1; Biocare), CD163 (clone 10D6; Thermo Fisher Scientific), CD4 (clone EPR6855; Abcam).

### Confocal Microscopy

Lungs were excised and submerged in BD Cytofix diluted 1:3 with PBS for 24hr at 4°C. Lungs were washed twice in PBS and dehydrated in 30% sucrose for 24 hours prior to embedding in OCT and rapid freezing in a 2-methylbutane and dry ice slurry. A cryostat was used to generate 20um sections, which were stained overnight with fluorescently conjugated antibodies at room temperature and cover-slipped with Fluoromount G mounting media (SouthernBiotech). Images were acquired on a Leica SP8X confocal microscope.

### Histocytometry

Histocytometry analysis was performed as previously described, with minor modifications (Gerner et al., 2012). Briefly, multiparameter confocal images were first corrected for fluorophore spillover using the built-in Leica Channel Dye Separation module. For acquisition of single stained controls, UltraComp beads (Affymetrix) were incubated with fluorescently conjugated antibodies, mounted on slides, and imaged. Cell surfaces were created using Jojo1 nuclear staining using the Imaris surface creation module, and the object statistics were exported to Excel (Microsoft). Object statistics were combined into unified CSV files and finally imported into FlowJo software for cellular gating and analysis. For correlation analyses, histocytometry data are spatially subdivided into 80um radius circular neighborhoods via virtual raster scanning. Pearson correlation coefficients are then calculated for the numbers of cells in these neighborhoods. For visual clarity, presented images were manipulated in Imaris and PowerPoint (Microsoft), with identical manipulation applied across experimental groups.

### Statistical analysis

Statistical tests were selected based on appropriate assumptions with respect to data distribution and variance characteristics. No statistical methods were used to predetermine sample size. The statistical significance of differences in mean values was analyzed by a two-tailed unpaired Student’s t test. Paired t test was performed only when comparing responses within the same experimental animal or tissue (indicated in the legend). Correlations and corresponding P values by Pearson’s correlation test. ****,p ≤ 0.0001; ***, p ≤ 0.001; **,p ≤ 0.01; and *, p ≤ 0.05; NS, p > 0.05.

### Lead Contact and Materials Availability

Further information and requests for resources and reagents should be directed to and will be fulfilled by the Lead Contact, Kevin Urdahl. This study did not generate new unique reagents.

## Supplemental Information titles and legends

**Figure S1. Related to Figure 1. Despite antigen sensing, CD4 T cells produce minimal IFNγ within Mtb granulomas:** A) Example histocytometry gating scheme of indicated cell populations within infected lungs B) Example segmentation of histocytometry data. 80um radius neighborhoods are generated using raster scanning as shown. C) Flow cytometry gating strategy for identification of Mtb-specific and OVA-specific T cells using congenic marker staining, as well as demonstration of IFNγ staining after ex-vivo restimulation. D) P-values from correlation analysis shown in Fig 1E. E) Paired flow cytometry data from figure 2, demonstrating ex-vivo IFNγ staining in Mtb-specific and control OVA-specific T cells. F) Analysis of relative Mtb-specific vs. OVA-specific T cell abundance as determined by flow cytometry vs. histocytometry. G) Analysis of IFNγ production by the peptide-stimulated, Th1 polarized CD4 T cells prior to adoptive transfer. Correlations and corresponding P values (D) by Pearson’s correlation test. Single-group comparisons by unpaired t test (F) or paired t test (E). Data are presented as mean ± SD. *p < 0.05, ***p < 0.001, ****p < 0.0001.

**Figure S2. Related to Figure 2. Genes associated with TGFβ signaling are increased within the granuloma:** A) GSEA analysis of parenchymal Mtb-specific cells vs. naïve T cells, showing increased pathways consistent with activation. B) GSEA analysis highlighting the indicated pathways which were differentially associated with the parenchymal vs. intravascular Mtb-specific cells. C) Relative expression of genes in the TGFβ signaling pathway in parenchymal vs. intravascular Mtb-specific cells is shown. D) Representative confocal image of a necrotic NHP granuloma, showing pSMAD3 staining. E) Zoom-in images showing pSMAD3 signal in relation to CD4 and CD68/163. FDRs were calculated using the Benjamini-Hochberg method.

**Figure S3. Related to Figure 3. Lack of TGFβRII on T cells results in decreased CFUs and increased IFNγ production:** A) Example flow cytometry gating strategy for the identification of parenchymal Mtb-specific T cells, as well as for the quantification of IFNγ production by these cells is shown. B) P-values from the correlation analysis shown in Fig 3G. C) Representative confocal images demonstrating staining of iNOS within CD11b+ cells. Correlations and corresponding P values (C) by Pearson’s correlation test.

**Figure S4. Related to Figure 4. TGFβ inhibits effector function of Th1 cells:** A) Flow cytometry gating strategy for identifying and phenotyping ESAT6 tetramer-positive TGFβR.KO vs WT cells in the lung parenchyma. B) Analysis of IFNγ MFI on IFNγ-expressing tetramer-positive WT vs TGFβR.KO cell cells is shown. Related to Main Figure 4A. C) Analysis of IFNγ production by the peptide-stimulated, Th1 polarized Tg.WT vs Tg.KO CD4 T cells prior to adoptive transfer. Single-group comparisons by paired t test. Data are presented as mean ± SD. *p < 0.05, ***p < 0.001, ****p < 0.0001.

