## Supplementary figures and images for "TGFβ restricts T cell function and bacterial control within the tuberculous granuloma"

### Supplemental Figure 1

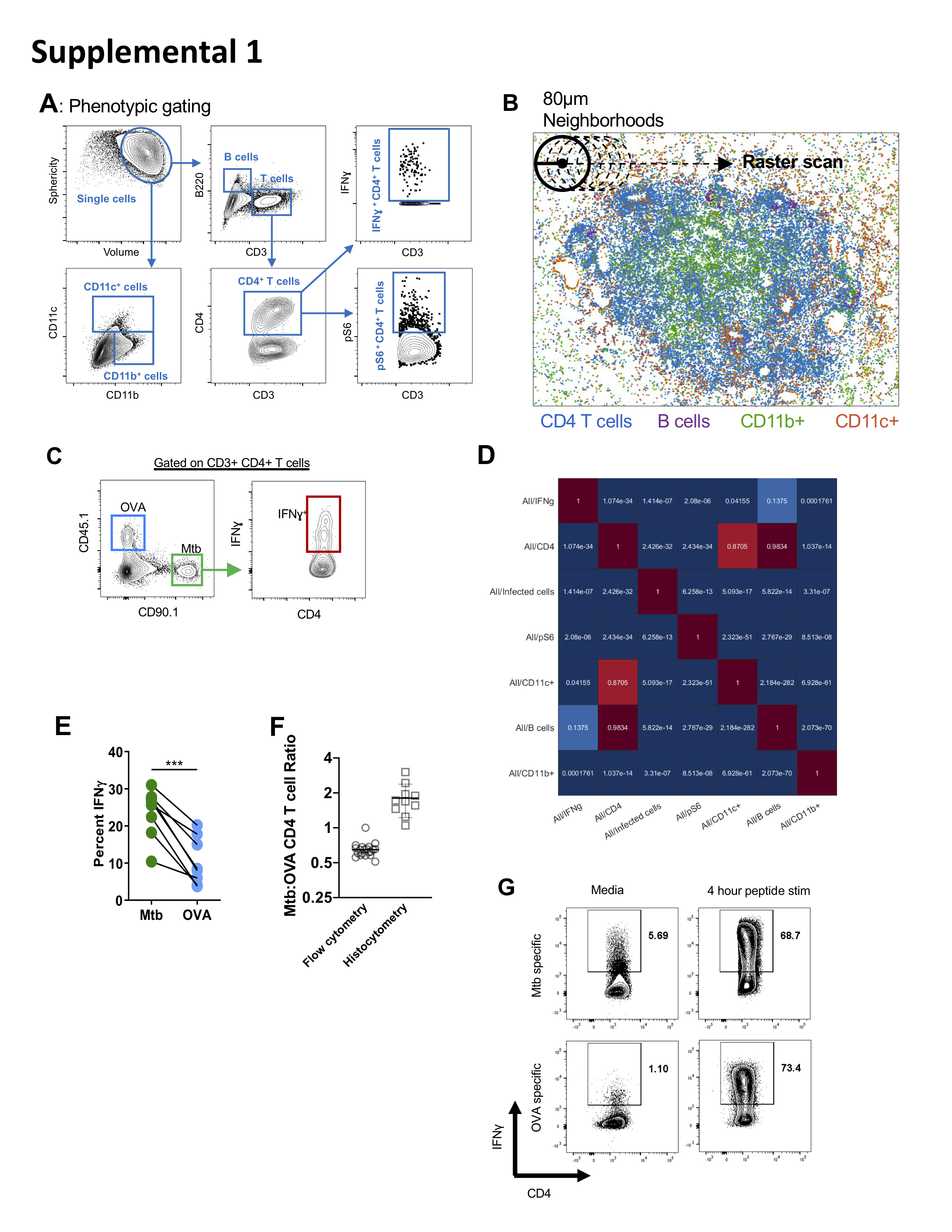

### Supplemental Figure 2

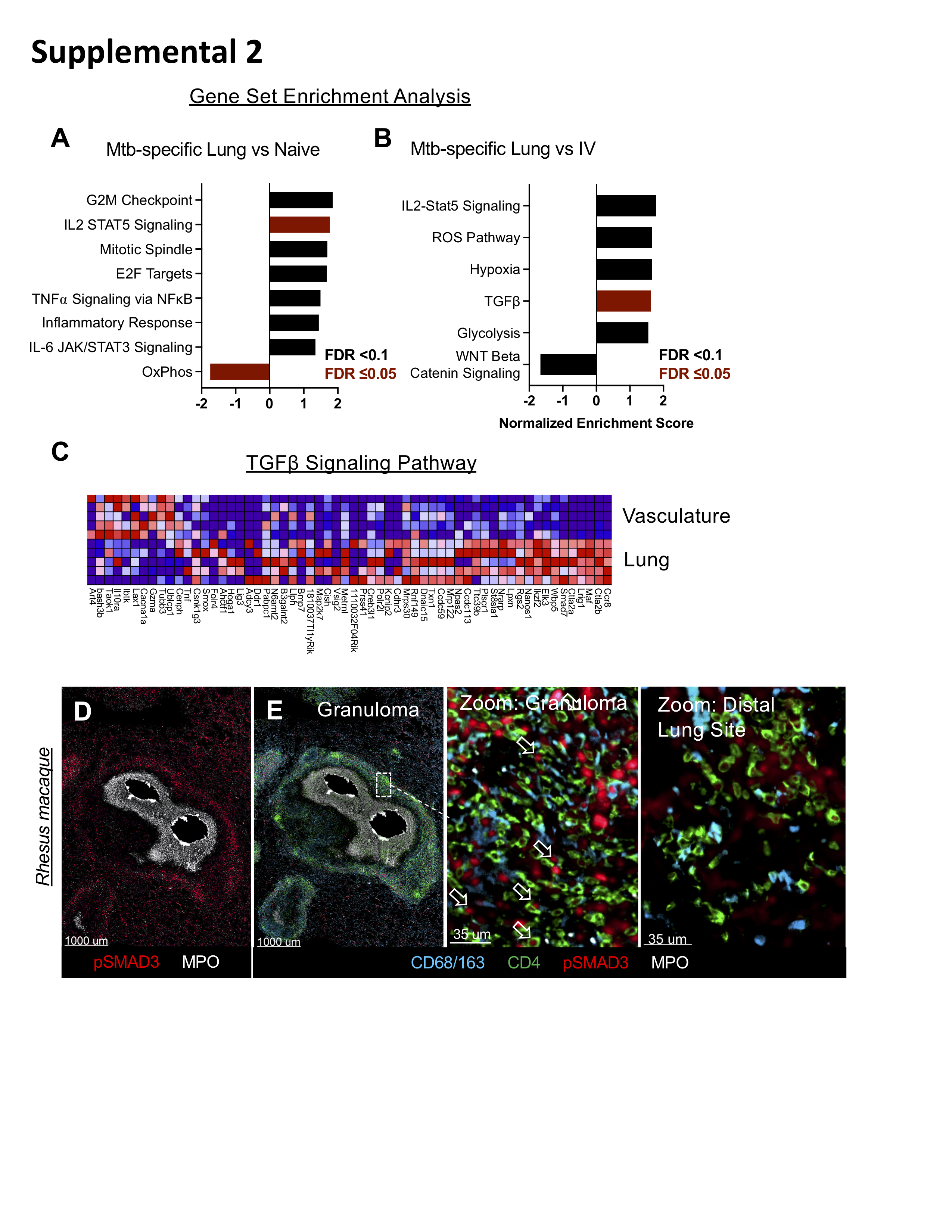

### Supplemental Figure 3

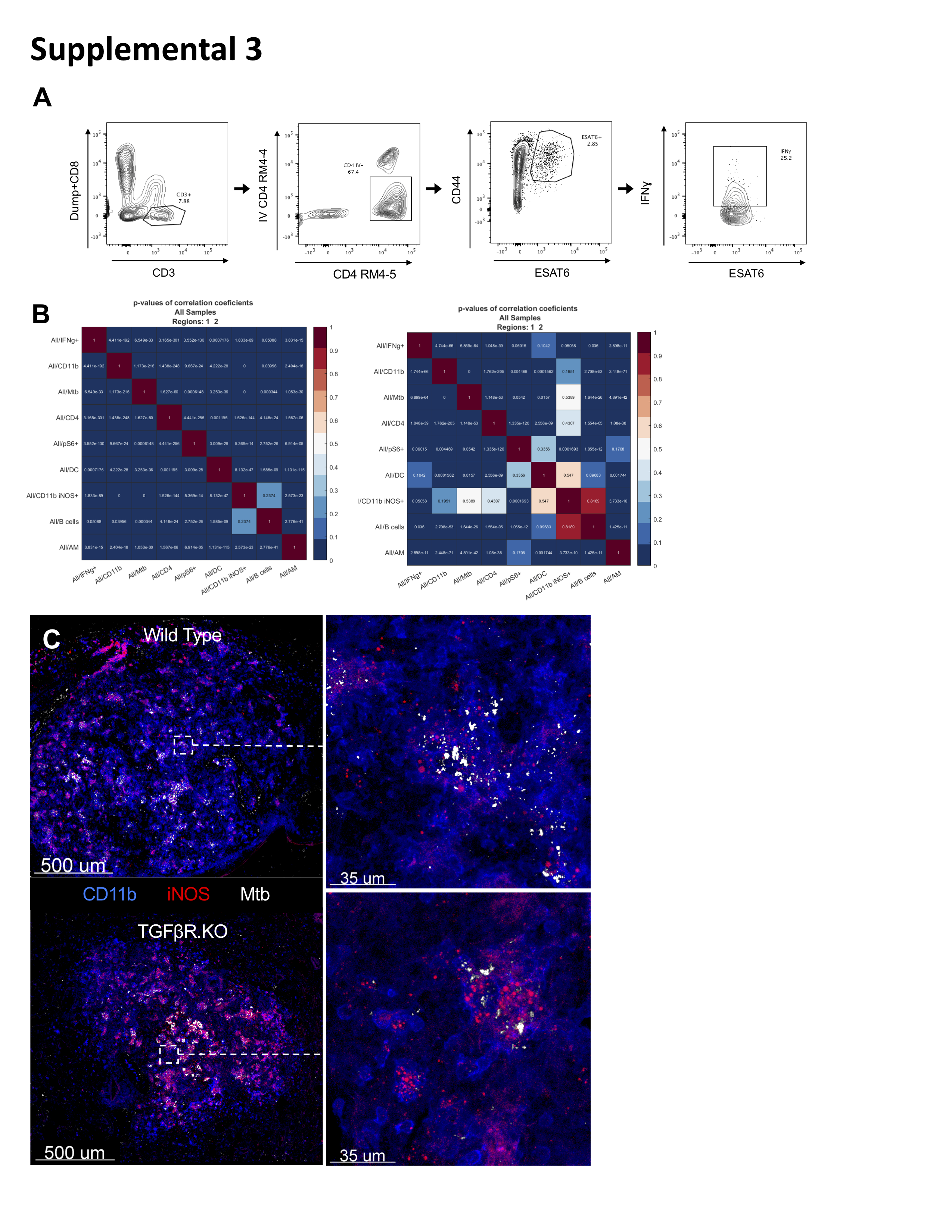

### Supplemental Figure 4

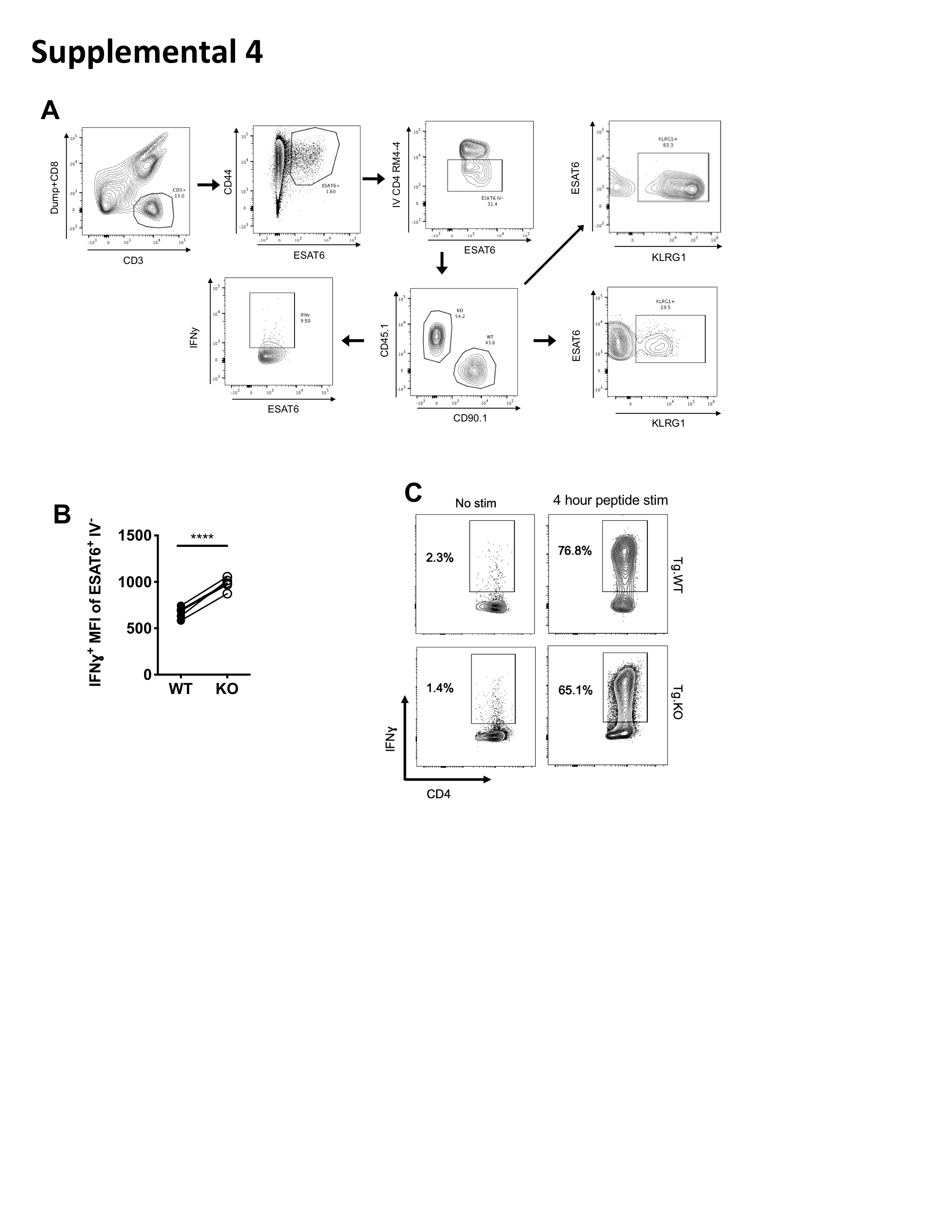
